# Genomics of a killifish from the Seychelles islands supports transoceanic island colonization and reveals relaxed selection of developmental genes

**DOI:** 10.1101/2020.08.03.232421

**Authors:** Rongfeng Cui, Alexandra M Tyers, Zahabiya Juzar Malubhoy, Sadie Wisotsky, Stefano Valdesalici, Elvina Henriette, Sergei L Kosakovsky Pond, Dario Riccardo Valenzano

## Abstract

How freshwater fish colonize remote islands remains an evolutionary puzzle. Tectonic drift and trans-oceanic dispersal models have been proposed as possible alternative mechanisms. Integrating dating of known tectonic events with population genetics and experimental test of salinity tolerance in the Seychelles islands golden panchax (*Pachypanchax playfairii*), we found support for trans-oceanic dispersal being the most likely scenario. At the macroevolutionary scale, the non-annual killifish golden panchax shows stronger genome-wide purifying selection compared to annual killifishes from continental Africa. Reconstructing past demographies in isolated golden panchax populations provides support for decline in effective population size, which could have allowed slightly deleterious mutations to segregate in the population. Unlike annual killifishes, where relaxed selection preferentially targets aging-related genes, relaxation of purifying selection in golden panchax affects genes involved in developmental processes, including fgf10.

## Introduction

Taxa with limited dispersal capabilities, such as flightless mammals, amphibians and freshwater fishes are thought to speciate largely through vicariance, i.e. population isolation caused by the presence of physical barriers, such as rivers and mountain ranges (Wiley 1988). Whether species with limited dispersal capacity, such as freshwater fishes, colonize remote islands through tectonic drift or trans-oceanic events is still debated (Briggs 2003; Sparks and Smith 2005).

Aplocheiloid killifish (Aplocheiloidei, Cyprinotondiformes) are small fresh-water and brackish water fishes distributed in South-east Asia, America, Africa, Madagascar, Seychelles and India. Killifishes are often adapted to survive in extreme environments, such as those characterized by seasonal drought. Some killifishes evolved an annual life cycle (**Figure 1**), which allows fertilized eggs to remain in diapause and survive through the dry season (Cellerino, et al. 2016). The main mechanism explaining the current distribution of killifishes across the planet is the mesozoic divergence of the major killifish clades, which followed the tectonic drift order of Gondwana (Murphy and Collier 1997; Costa 2013). Hence, if taxa mainly speciated as a consequence of geographical isolation due to tectonic events, the phylogenetic branching order and timings are expected to agree with those of continental drift.

**Figure 1.**
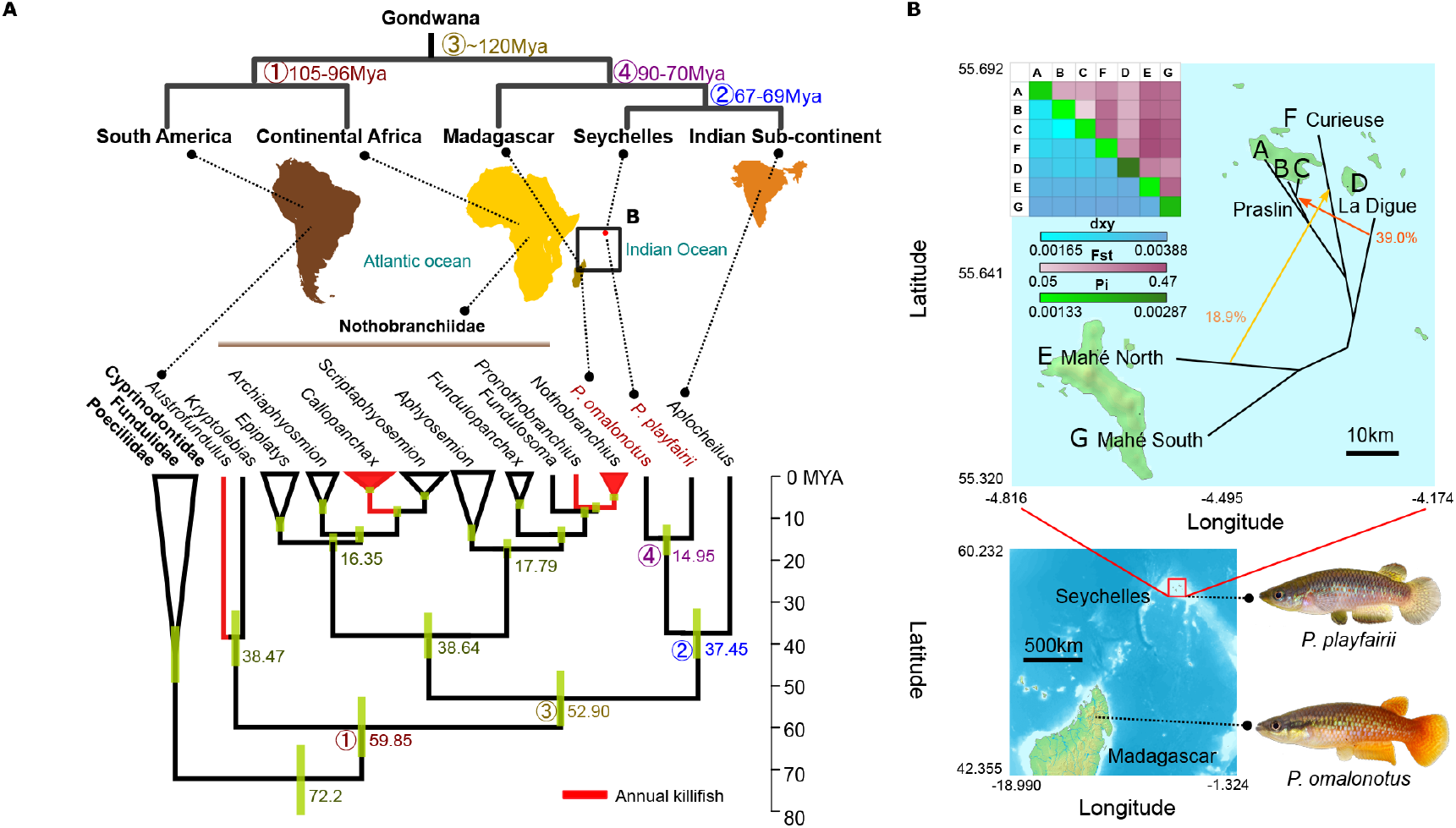
**A.** Dated phylogeny of Cyprinodontiforms mapped to tectonic events of the Gondwana continents. Tree was built with RAxML with 100 rapid bootstraps, and MCMCtree with GTR+LogNormal clock was used to infer divergence times, calibrated from a previous teleost phylogenetic study (external taxa trimmed off in figure). Node bars showing the 95% C.I. from the posterior distribution of divergence times. The order of continental breakup and their timings do not fit the inferred divergence times of major killifish clades. Putative tectonic event-driven nodes are numbered 1 through 4, with the mean divergence times indicated. **B.** Distribution of *P. omalonotus* and *P. playfairii* on Madagascar and the Seychelles. Population history of sampled populations of *P. playfairii* was inferred by TreeMix, allowing for two migration edges colored orange (weight given in percentages).

However, the genus *Pachypanchax* challenges this model. *Pachypanchax* are non-annual killifishes restricted to Madagascar and the neighboring Seychelles, with the sister genus *Aplocheilus* found in India and South-east Asia. Only a single species, *Pachypanchax playfairii* (the golden panchax) is found on the Seychelles islands. Despite the current geographical proximity to Madagascar, the Seychelles islands have a closer geological relationship with the Indian subcontinent (Plummer and Belle 1995; Seton, et al. 2012). Hence, based on the continental drift hypothesis of taxa distribution, species from Seychelles with limited dispersal capacity, like freshwater fish, are expected to have closer phylogenetic proximity to India than to Madagascar. However, Seychelles and Madagascar are home to fish of the same genus (*Pachypanchax*), suggesting the possibility for transoceanic dispersal following tectonic drift as a possible mechanism for species dispersal. Evidences for transoceanic dispersal have been found in other groups of animals. For example, palaeontological and molecular clock analyses place the divergence time of cichlid fish in the Palaeocene (65-57 Myr ago), much later than Mesozoic tectonic rifts of Gondwana (Friedman, et al. 2013). Based on molecular clock and nucleotide substitution saturation analysis in mitochondrial genes, Malagasy amphibian, reptilian and non-flying mammals are too closely related to continental sister groups to fit the ancient tectonic events (Vences 2004). The Malagasy chameleon *Archaius tigris* was shown to be sister to a continental African chameleon genus, which diverged not until the Eocene–Oligocene, again supporting the possibility for transoceanic dispersal (Townsend, et al. 2011). Molecular dating of the now extinct Malagasy Elephant Bird reveals its relatively recent divergence with the sister group New Zealand Kiwis, suggesting transoceanic dispersal by their flying ancestors (Grealy, et al. 2017).

To directly test whether tectonic or transoceanic dispersal can explain the evolution of the golden panchax (*Pachypanchax playfairii*), we used phylogenetic and population genomic methods, as well as a lab experiment. The golden panchax has a long evolutionary history of isolation on the Seychelles islands lasting several million years, which combined with its limited dispersal capacity, represents a paradigmatic example of island evolution. Evolution through island colonization has been used as an idealized model for understanding speciation (MacArthur and Wilson 2001). Endemic species are known to be more susceptible to extinction attributed to inbreeding (Frankham 1998), despite some taxa evolving mechanisms for its mitigation (Wheelwright and Mauck 1998). Reduction of effective population normally lowers the efficacy of selection, leading to the accumulation of deleterious mutations (Charlesworth, et al. 1993). On the other hand, shifts in the ecological niche can also lead to relaxed selection by removing niche-specific selective constraints (Wertheim, et al. 2014). Colonizing new island habitats is likely accompanied by both a reduction of effective population size and by a change of the ecological niche, which may affect a suite of phenotypes. For instance, relaxation of pre-existing selective pressures, such as a lack of predators in novel environments, can explain the loss of anti-predator behaviors in oceanic tortoises (Austin and Nicholas Arnold 2001) and the loss of flight in the Kakapo (Livezey 1992) and Dodo (Livezey 1993). Positive selection has also been invoked to explain island-specific phenotypes. For example, gigantism in oceanic tortoises is proposed to confer a selective advantage during the unpredictable long dry seasons on islands (Jaffe, et al. 2011). However, still little is known about the genomic patterns of species undergoing long-term island isolation.

Previous studies conducted in continental African killifishes have shown that relaxation of selection dominates genome evolution in short-lived annual African killifishes, preferentially targeting aging-related pathways, likely resulting from population fragmentation, population size reduction and a shift in the selective regime (Cui, et al. 2019; Willemsen, et al. 2019). Here, to assess the effects of island isolation in the golden panchax, we inferred the historical changes in effective population size in wild populations from different islands of the Seychelles archipelago and conducted an unbiased characterization of the relative contribution of relaxation of selection, intensified selection and positive selection in shaping the golden panchax genome.

Our results show that golden panchax likely colonized the Seychelles via transoceanic dispersal and indicate that prolonged insularism led to progressive population size reduction. Detectable relaxation of purifying selection in golden panchax appears to largely affect developmental genes, including fgf10 (important for cell cycle and organ development), and is markedly distinct from the pattern observed in annual killifishes from continental Africa.

## Results

### The killifish biogeographic pattern supports transoceanic species dispersal

We constructed a phylogenetic tree using 4026 orthologous protein-coding sequences from 61 teleosts, adding the newly sequenced genome of the Madagascar species *Pachypanchax omalonotus* (estimated genome size ~ 707 Mbp). Based on calibrations obtained from a broader-scale teleost phylogeny (Near, et al. 2012), we inferred divergence times with mcmctree (**Figure 1A**). South American representatives of the family Rivulidae diverged from African and Indo-malagasy families ~59.85 myr ago (95% CI 52.66-67.08). Nothobranchiidae, which are found in continental Africa, diverged from the Indo-malagasy Aplocheilidae ~52.90 myr ago (95% CI 46.33-59.66). Continental African killifishes are deeply diverged into western and eastern clades ~38.64 myr ago (95% CI 33.18-44.07). The Indian genus *Aplocheilus* diverged from *Pachypanchax* ~37.45 myr ago (95% CI 31.55-43.50). Amongst all sampled genera with more than one representative species, *Pachypanchax* has the most ancient common ancestor at ~14.95 myr ago (95% CI 11.47-18.77), even deeper than the divergence between the sister genera *Nothobranchius* and *Pronothobranchius*, which dates to ~8 myr ago (95% CI 6.87-9.22) and between *Callopanchax* and *Scriptaphyosemion*, which dates to ~8.9 myr ago (95% CI 7.61-10.32)(**Figure 1A**). The divergence between the two extant species in the genus *Pachypanchax* is comparable with the divergence between more distantly related genera, such as *Fundulopanchax* and *Nothobranchius* (14.4 myr ago, 95% CI 12.59-16.50) or *Archiaphyosemion* and *Callopanchax* (14.5 myr ago, 95% CI 12.45-16.41). To our surprise, neither the order of divergence nor any of the major divergence dates fit the known tectonic order of the Gondwanan continents, suggesting that transoceanic dispersal rather than tectonic drift may explain the current biogeographic pattern within killifish (**Figure 1A**). We also found that although the concatenated mitochondrial tree using codon positions 1 & 2 has only a moderate rapid bootstrap support of 69 and a non-significant AU test statistics of 0.264 (Shimodaira 2002), the topology of the maximum likelihood tree agrees with the nuclear genome (**Figure S1**), a result contradicting previous mitochondrial trees built with fewer genes (Murphy and Collier 1997). Therefore, we find that the tree topology from mitochondrial and nuclear genetic data are largely in agreement across all the major branches of killifish and support a significant contribution of transoceanic species dispersal to the current killifish species distribution.

Studying population separation among golden panchax (*Pachypanchax playfairii*) from the Seychelles reveals a branching pattern consistent with island distance (TreeMix analysis, **Figure 1B**); and the genome-wide statistics of divergence *dxy* and *Fst* agree with the migration analysis based on TreeMix tree (**Figure 1B**). Overall, the level of divergence between populations has the same range as within-population polymorphisms, suggesting recent divergence. Interestingly, samples from northern Mahé island are very divergent from samples from the south, but we cannot rule out the difference being driven by introduction of domesticated fish samples of the Mahé North island. TreeMix analysis supports adding one migration edge from the La Digue population to the (B+C) population on the neighboring Praslin island, with an estimated edge weight of 39-41%, suggesting high gene flow (**Figure 1B**). The resulting divergence among populations remains unchanged with outgroup inclusion (**Figure 2**). Addition of further migration edges is not supported as even when one migration was added, the network already explains on average 99.5% (outgroup excluded) or 99.99% (outgroup included) of the variation in the data (**Figure S2**).

Together, our results suggest that both macro- and micro-evolutionary divergence of killifishes can be in part explained by transoceanic dispersal.

**Figure 2.**
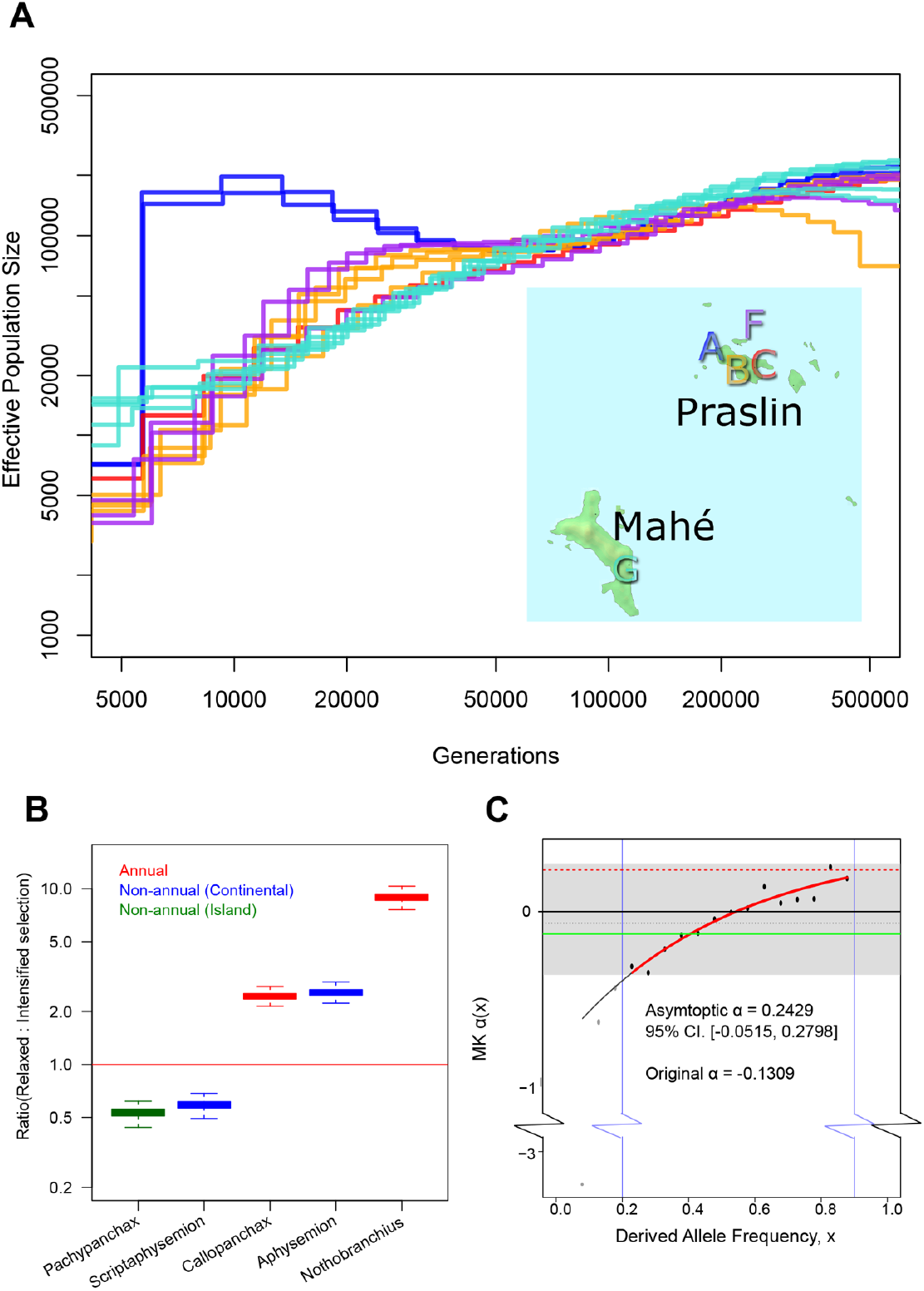
**A.** PSMC plot for wild-caught *P. playfairii* individuals from 3 localities in Praslin, one from Curieuse and one from Mahé. **B.** Ratios plotted on a logarithmic scale of gene counts that are detected as relaxed and intensified selection in five killifish genera. Relaxed selection detected by the 2-k parameterization, intensified selection with the original 1-k parameterization. Confidence interval obtained by 1000 multinomial resampling. Red line indicates ratio at 1. **C.** McDonald-Kreitman α plotted against derived allele frequencies, outgroup *P. omalonotus*.

### *Pachypanchax* embryos develop normally in sea water

To assess whether golden panchax would survive extended periods of time in the sea, which would be consistent with the transoceanic species dispersal supported by the genomic data, we incubated in seawater golden panchax embryos and compared their developmental trajectory to embryos raised in normal conditions, i.e. freshwater incubation. Remarkably, we found that golden panchax embryos develop perfectly in seawater, with no detectable differences in survival with control groups incubated in freshwater (**Figure S3**). Moreover, fry from golden panchax embryos raised in seawater remain viable. This finding does not extend to the continental African killifish *Nothobranchius furzeri*, whose embryos did not survive in seawater, with all embryos dead after 3 days post collection. Our findings suggest that golden panchax could in principle sustain extended periods of time in seawater, which could be compatible with a model of direct transoceanic dispersal vial ocean currents.

### Annual killifishes from Madagascar and Seychelles accumulate deleterious mutations as population size declines

Annual killifish from continental Africa display a larger proportion of genes under relaxed rather than intensified selection compared to related non-annual species (Cui, et al. 2019). We asked whether the two annual species from Madagascar and Seychelles (i.e. *Pachypanchax omalonotus* and *playfairii*), followed the same pattern of genes under relaxed vs. intensified selection observed in the annual killifish species from continental Africa. We used phylogenetic methods to examine intensity of selection at the macroevolutionary scale. We identified 444 genes under relaxed purifying selection (p < 0.05, **Table S4**) and 877 genes under intensified selection (p < 0.05, **Table S5**) in the genus *Pachypanchax* relative to other aplocheiloids. Compared to genera of annual killifish within the Nothobranchiidae family, the genus *Pachypanchax* has a larger proportion of intensified rather than relaxed genes (**Figure 2B, Figure 3**), at a similar level as the non-annual killifish genus *Scriptaphyosemion* (from continental Africa). We used a previously simulated dataset to compare the performance of the one-scaler and two-scaler parameterizations of the RELAX test (Cui, et al. 2019). The simulated dataset was based on parameters obtained from the non-annual *Scriptaphyosemion* clade. We simulated genes that are under five selection regimes: relaxed purifying selection, intensified purifying selection, relaxed and positive selection, intensified and positive selection, and positive selection only. Then we ran both the RELAX tests on these datasets and recorded the number of genes called as relaxed or intensified. We found that the two-scaler parameterization of RELAX has 7.6% higher power to detect relaxed selection when positive selection is also present in the same gene (**Table S2**). Simulations also show that the original RELAX parameterization has a higher power (46.7% more) for detecting intensified selection (**Table S2**); we therefore used this method for intensified selection, after excluding any genes already called relaxed by the two-scaler parametrization method. Overall, our findings show that the island annual species of the genus *Pachypanchax* have a larger proportion of genes under intensified rather than relaxed selection compared to annual killifishes, to a level that is comparable to non-annual killifish genera living in continental Africa. Nevertheless, the total number of genes (444) under relaxed selection in *Pachypanchax* is still much higher than the simulated false positive rate (**Table S3**; expected = 253-269 in 10556, assuming 1-10% intensified genes, 90-99% genes with no shifts in selection and 2.5% positively selected genes) of the test, suggesting that relaxed selection still fixed deleterious mutations in this non-annual genus.

**Figure 3.**
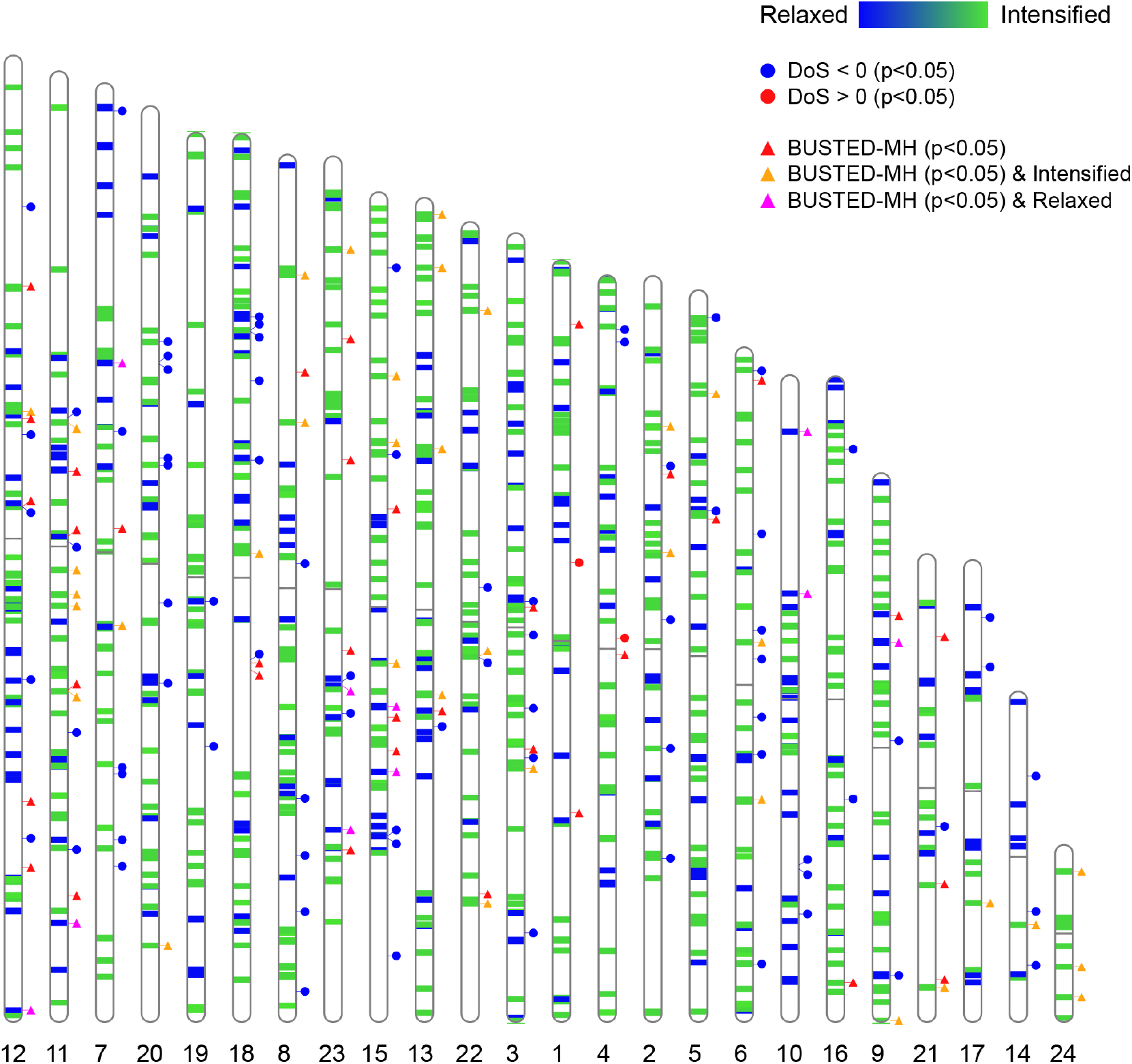
Ideograms showing genes under relaxed purifying selection (dark blue) and intensified selection (bright green) in the *Pachypanchax* clade plotted on 24 pseudochromosomes. Relaxed selection detected with HYPHY RELAX with 2-k parameterization, intensified selection detected with the original 1-k parameterization. Genes under positive selection (triangles) detected in the *Pachypanchax* lineage, accounting for multinucleotide substitutions and mutational rate variation using HYPHY BUSTED-MH. Direction of Selection (DoS) statistics computed for wild-caught *P. playfairii* individuals from Praslin, La Digue and Curieuse, using *P. omalonotus* as the outgroup.

To investigate the possible cause of relaxation of selection in *Pachypanchax*, we focused on the more isolated Seychelles species golden panchax (*Pachypanchax playfairii*). We collected individual golden panchax from several natural localities (**Table S1**), sequenced their genomes, and we conducted population genetics analysis, using *Pachypanchax omalonotus* as outgroup. We pooled individual fish sequences from the Praslin, Curieuse and La Digue to analyze the asymptotic McDonald-Kreitman alpha (Messer and Petrov 2013). A positive alpha value is an estimate of the proportion of substitutions fixed by positive selection since divergence from *P. omalonotus*. Negative alpha is due to the segregation of slightly deleterious mutations in *P. playfairii* since divergence from the outgroup (Messer and Petrov 2012). The estimated ratio of mutations fixed by positive selection is not significantly different from zero, suggesting not many sites were positively selected in this Seychelles taxon. At lower derived allele frequencies, the alpha statistics remains negative and does not reach the asymptotic level until 80% of derived allele frequency, indicating that slightly deleterious variants are segregating at high frequency in the population (**Figure 2C**). Therefore, although the macroevolutionary trend suggests that the *Pachypanchax* clade has more intensified selection compared to other killifishes, the MK alpha measurements indicate the presence of some slightly deleterious variants. We then asked what could be the leading cause for the extensive number of genes under relaxed selection and of the deleterious gene variants segregating at high frequency in the golden panchax. A likely scenario is that decreased effective population size may have occurred in this island species. To test this hypothesis, we then analyzed individual genomes from wild-caught golden panchax using PSMC’, and found that most populations underwent a gradual decline in effective population size since 200,000 generations ago, ending in a population size of about 5000-20,000 at generation 5000 (**Figure 2A**). Different individuals from the same population show repeatable PSMC’ traces. Population A from the Praslin island had a population expansion from ~20,000 generations ago until ~7,000 generation ago, after which it rapidly declined to less than 10,000. Overall, island annual killifish display a larger proportion of the genome under intensified rather than relaxed selection, compared to annual killifishes from continental Africa. However, their extended isolation and progressive reduction in population size may have caused the genome-wide accumulation of nearly neutral polymorphisms at otherwise conserved sites in more recent generations.

### Relaxation of selection in non-coding regions

Besides protein-coding genes, molecular evolution of regulatory elements of genomes underlies important phenotypic variations in response to selection in different organisms, including the loss of flight in Ratite birds (Sackton, et al. 2019) and degeneration of armor plates in Stickleback fish (O'Brown, et al. 2015). We therefore explored whether non-coding regions in the *Pachypanchax* genome experienced relaxation of selection. First, we identified 50bp conserved non-coding elements (totaled 86,087,429bp, excluding CDS 56,842,251 bp, excluding repeats 54,779,098bp) in the assembled genomes of the annual species *Austrofundulus limnaeus*, *Callopanchax toddi*, *Nothobranchius furzeri* and *Nothobranchius orthonotus* using PhastCon. These elements are expected to have highly conserved regulatory functions within a ~60myr divergence time. Together, 10223 (511.15kb, or 0.093%) of these elements were significantly accelerated (i.e. had significant sequence divergence compared to the annual species) in the *Pachypanchax* genome when compared to 4-fold degenerate sites. Because 4-fold degenerate sites are known to evolve slower than completely neutral sites due to codon bias (Hershberg and Petrov 2008), we interpret these accelerated elements to have likely undergone relaxed purifying selection in the genus *Pachypanchax*. We then asked whether these accelerated conserved elements were enriched for regulatory sequence motifs (using DREME), and found enrichment for the promoter TATA box (TAATTA, p= 5.6e-28) and the enhancer box (CAGCTG, p = 6.5e-13), corroborating their potential role as regulators of gene expression. Then, we asked whether these non-coding accelerated conserved elements were associated with downstream genes that evolved under intensified or relaxed selection. We found that protein-coding genes within 80kb distance from an accelerated conserved non-coding element were more likely to be also under relaxed selection, according to the 2-scaler RELAX test. Furthermore, the median direction of selection (DoS, where negative values are indicative of genes evolving under relaxed selection) was more negative for protein-coding genes near an accelerated conserved non-coding element (**Figure S4**). Overall, we found that relaxation of selection in *Pachypanchax* extends beyond the coding regions, supporting a scenario where both gene regulation and adjacent coding sequences are subject to similar fluctuations of evolutionary constraints.

### Intensified selection on RNA polymerase complex genes

We performed a GO term enrichment analysis on intensified genes (i.e. under positive or purifying selection) detected with the original RELAX method (more powerful in identifying genes under intensified selection, based on our simulations), after excluding genes detected to be relaxed by the new parameterization (more powerful in identifying genes under relaxed selection, based on our simulations). Eight biological complexes were found to be enriched at an FDR cutoff of 0.05. The most significantly enriched component was the RNA polymerase complex (p = 2.12e-6, FDR = 0.000338), containing 19 genes detected to have undergone intensified selection in the *Pachypanchax* branch, including 6 RNA polymerase subunits and 4 general transcription factor subunits. Other complexes under intensified selection (FDR < 0.05) include the MICOS complex, smooth ER, nuclear envelope lumen and transferase complex (**Table S6**).

### Relaxed selection affects genes involved in early *Pachypanchax* development

Despite having some portion of the genome under relaxed purifying selection, fish of the genus *Pachypanchax* live several years, which is a significantly longer time than the corresponding annual species from continental Africa, whose lifespan can be as short as 4 to 6 months (Valenzano, et al. 2015; Willemsen, et al. 2019). Since relaxation of purifying selection largely affects genes involved in late-life maintenance in annual African killifishes (Cui, et al. 2019), we asked what genes and gene functions were affected by relaxed selection in a clade with long-term island isolation yet maintaining a normal lifespan. To answer this question, we performed gene ontology analysis on the genes under relaxed selection (**Figure S5**; **Table S7**). Terms related to early development, such as anatomical structure morphogenesis (p=7.00e-5, FDR = 0.0012), cell communication (p=1.47e-6, FDR=5.19e-5) and cell surface receptor signaling pathway (p=1.68e-7, FDR=6.51e-5) are strongly enriched in genes under relaxed selection. Further pathway analysis using ConsensusPathDB (Herwig, et al. 2016) of the same gene set reveals that 7 pathways are enriched (**Table S8**; FDR < 0.05). Among these is the fibroblast receptor 2 ligand-binding pathway (p=0.000495, FDR=0.046), which has been found to be important for early embryogenesis. Aligning the protein sequences of human and killifishes, we found that both *Pachypanchax omalonotus* and *Pachypanchax playfairii* carried a serine instead of a glycine at a critical position on fgf10, while this position is conserved from human to other fishes (**Figure 4**). This amino acid is homologous to G160 in human fgf10, which in turn interacts with R251 on the human receptor fgfr2 (Yeh, et al. 2003). The homologous amino acid R251 of fgfr2 is conserved in all killifishes, including *Pachypanchax*.

**Figure 4.**
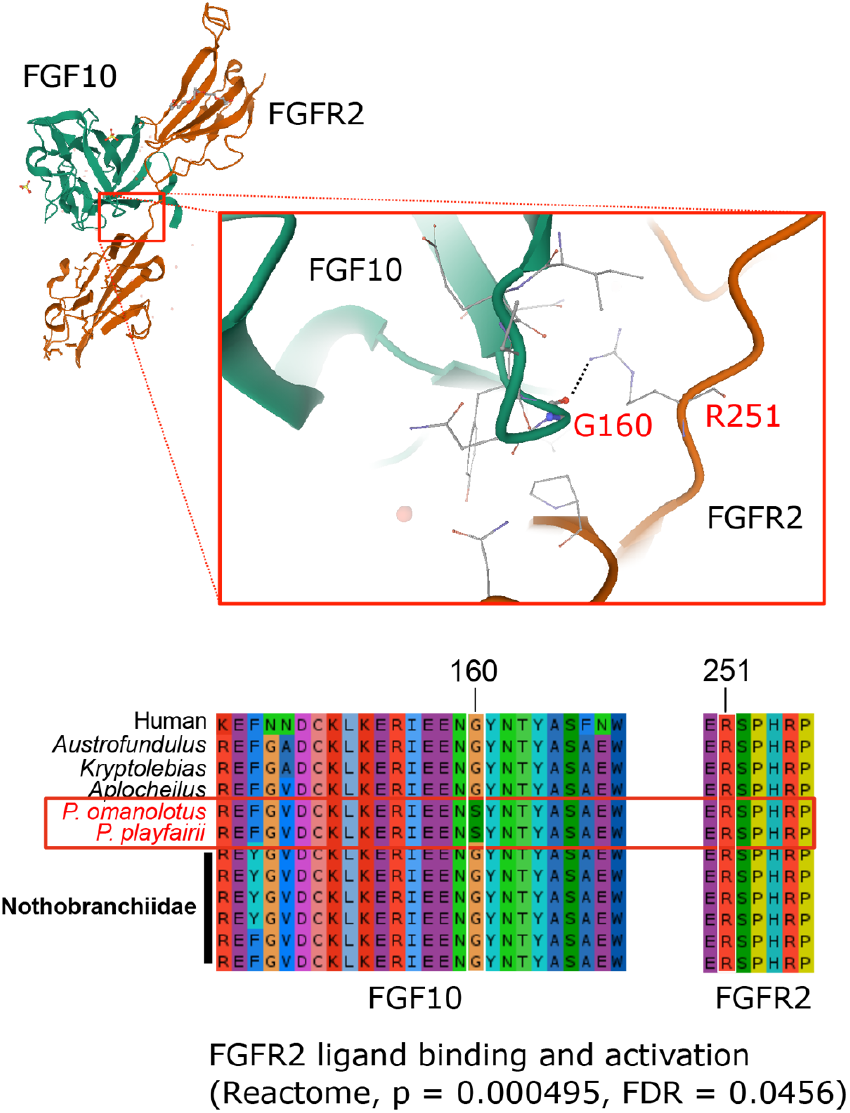
Structure of the human FGF10-FGFR2 complex, focusing on the interaction of G160 and R251 (Yeh, et al. 2003). Partial protein sequence alignments of FGF10a and FGFR2 showing key interacting residues G160 and R251. Two *Pachypanchax* species have a unique substitution from Glysine to Serine corresponding to residue 160 in human FGF10. The key amino acid at position 251 corresponding to human FGFR2 is conserved.

Since we found that relaxation of selection applies to both coding and non-coding regions, we asked what genes and gene functions were enriched in proximity of accelerated non-coding elements in the genus *Pachypanchax*. This analysis confirmed that accelerated non-coding elements map in proximity of genes involved in developmental processes (**Table S9**). Therefore, not only did we find a strong correlation between relaxation of selection in coding and adjacent non-coding regions (**Figure S4**), but both of the GO enrichments also support that non-annual fish of the genus *Pachypanchax* underwent relaxation of selection at specific developmental genes.

### Minimal evidence of positive selection in *Pachypanchax*

To detect positive selection in the genus *Pachypanchax*, we ran BUSTED-MH (**Table S9**), confirming that when correcting for multi-nucleotides substitution and mutation rate variation, there is little evidence of positive selection in this clade, as less than 5% of tested genes were called as under positive selection, not exceeding the p value cutoff of 0.05 (**Figure 3**). Gene ontology analysis of candidate positively selected genes reveals no enriched terms below an FDR cutoff of 0.05 (**Table S10**). Therefore, the higher proportion of genes under intensified rather than relaxed selection in *Pachypanchax* could be due to genes under purifying selection, rather than under positive selection.

## Discussion

### Transoceanic dispersal is compatible with killifish species distribution

There has been a great deal of debates on the extent to which tectonic events can explain the biogeographic pattern of Gondwanan fauna. Killifish distribution have been previously suggested as an example of speciation following the sequence of events that led to the separation of the Gondwanan plates, largely supported by the congruence between mitochondrial phylogeny and the order of break-up of the geological tectonic plates. However, a small-scale phylogenetic tree built on nuclear genomic information (Pohl, et al. 2015) has presented a different branching pattern, placing continental African killifishes as sister species to the Indo-Malagasy clade. Here we present a whole-genome phylogenetic tree, based on nuclear genome, which rejects the proposal of sister relationship between South American and continental African taxa. Instead, our results support that Indo-Malagasy and continental African taxa form a sister relationship. Furthermore, contrary to the species distribution predicted by the order of events congruous with tectonic drift, Seychelles *Pachypanchax* are closely related to Malagasy taxa, rather than to the Indian genus *Aplocheilus*. Molecular dating further shows that the 95% posterior distributions of divergence times are more recent than the proposed separation time of the corresponding continental plates. Therefore, a more likely explanation for the current distribution of killifishes must involve crossing oceanic barriers (**Figure S6**). However, the mechanisms by which transoceanic migration might have occurred is unknown. Open-ocean “rafts” and dispersal by birds have been suggested as plausible means (Silva, et al. 2019). In our work, we show that the freshwater golden panchax (*Pachypanchax playfairii*), a non-annual killifish from the Seychelles islands, can complete embryonic development and successfully hatch in seawater, providing a surprising support for the opportunity of transoceanic dispersal by means of direct embryo or juvenile transport via oceanic currents.

### Long term decline of population size accompanied by limited relaxed selection and positive selection

The amount of fixed and segregating deleterious mutations in *Nothobranchius* has been shown to negatively correlate with effective population size (Cui, et al. 2019; Willemsen, et al. 2019). Hence, we asked whether a similar trend occurred in non-annual species, such as the golden panchax (*Pachypanchax playfairii*). While population size steadily declined over the past 200,000 generations in most of the *golden panchax* populations we analyzed, the ratio of genes under relaxed selection detected with a phylogenetic method is lower than in genera within Nothobranchiidae, especially when compared to the annual genera. Testing deleterious mutations at a more recent time-frame using the McDonald-Kreitman alpha metrics, we found evidence of deleterious mutations being segregating at various frequencies, although probably not yet fixed, hence likely preventing their detection by the phylogenetic method (which scores only the more frequent or fixed alleles). Compared to populations of annual killifishes of the genus *Nothobranchius*, individuals from *Pachypanchax* populations have lower levels of polymorphism, and yet they fixed fewer deleterious mutations. This is not surprising, however, because accumulation of deleterious mutations is not only a function of population size, but also relates to the selection regime, which eventually affects the shape of the distribution of fitness effects (DFE) of new mutations. Golden panchax live in permanent water streams and are robust and long-lived fish, unlike *Nothobranchius*, which reside in ephemeral waters and are short-lived. Despite the small population size, natural selection acting over a long evolutionary time may have further prevented life-shortening mutations from accumulating in this clade.

In agreement with recent findings that positive selection is in general difficult to detect especially in the presence of repeated relaxed purifying selection (Chen, et al. 2020), we found weak evidence for positive selection in the golden panchax using both population genetics and phylogenetic methods.

The limited genetic diversity of the extant golden panchax populations, together with their limited distribution within the Seychelles archipelago and to their progressively declining effective population size, represents a serious concern for the future survival of this species, and calls for effective conservation measures aimed at long-term preservation of this species.

### Relaxed selection in *Pachypanchax* affects embryonic developmental gene pathways

Despite relatively fewer genes having undergone relaxation of purifying selection in *Pachypanchax* compared to annual killifishes, they still amount to hundreds. In annual killifish genera from continental Africa, genes under relaxed selection are related to longevity and are differentially expressed between young and old fish (Cui, et al. 2019). In contrast, genes under relaxed selection, as well as accelerated non-coding elements in *Pachypanchax* are significantly enriched for early embryo development genes. An intriguing scenario compatible with developmental genes evolving under relaxed selection in *Pachypanchax* is that relaxation of selection could occur in both regulatory regions and coding regions of genes related to embryo diapause. If diapause were an ancestral trait to both annual and non-annual killifishes, once the ancestor of *Pachypanchax* colonized permanent water streams, they might have lost the selective pressure to remove any deleterious mutation affecting diapause-related developmental genes. Interestingly, although we did not detect extensive positive selection or highly enriched gene ontology terms in candidate positively selected genes, the GO terms with a FDR cutoff from 0.05 to 0.2 for genes under positive selection are also related to developmental processes (cell cycle and semaphorin receptors). This could hint that remodeling of early developmental processes may have occurred in the ancestors of *Pachypanchax*, involving both relaxation and positive selection for the same gene pathways. We provide a list of positively selected sites as candidate targets for future studies in **Table S11**. Hence, by detecting relaxed selection in a non-annual killifish clade that lost embryonic diapause, we may help reveal the genetic architecture of this unique trait that characterizes annual killifishes.

## Conclusions

Salt water barriers represent a considerable limit to the diffusion of freshwater taxa. Our analysis of the branching pattern as well as our molecular dating results suggest that at least in Cyprinodontiforms, transoceanic dispersal offers a likely mechanism to explain the current biogeographic pattern, and is supported by specific physiologic adaptation (embryo development in seawater). We show that long-term isolation may lead to a steady decline in the effective population size, which can cause recent relaxed selection leading to deleterious variants segregating in the population. In permanent water, relaxed selection does not preferentially target aging related pathways. Instead, relaxed selection targets both protein-coding and non-coding regions related to embryonic development.

## Methods

### Sample collection

Specimens from most populations were collected during a dedicated field trip in April 2016 (and by Elvina Henriette on the following month), using dip nets. Fish were identified in the field, small fin clips were taken from their caudal fin and stored in 98% ethanol. Fish were then released back to their habitat. Voucher specimens (a random subsample of both sexes) were taken from most populations. All field sampling and export procedures followed regulations of Seychelles, with permits and research associateship issued by Seychelles Bureau of Standards Ref.A0157 dated 22 March 2016. Samples from Mahe North were from aquarium strains bred in captivity since 2013, for approximately 3-4 generations (**Table S1**).

### Library preparation and sequencing

Library preparation follows previous methods (Rowan, et al. 2015; Cui, et al. 2019). Briefly, genomic DNA was extracted by incubating a small finclip (1mm × 2mm) in 50 μL lysis buffer (Tris-HCl pH=8.0 10mM, EDTA 10mM, NaCl 10mM, SDS 0.5%) containing 0.5 μL Proteinase K (20mg/ml, NEB) at 50°C for 2hr. Forty μL SPRI beads (1mL SeraMag GE Healthcare, 65152105050250 beads in 100mL of PEG8000 20%, NaCl 2.5M, Tris-HCl pH=8.0 10mM, EDTA 1mM, Tween20 0.05%) were added directly to the mix to purify DNA with two washes by 80% ethanol. For library preparation, 250ng of genomic DNA was digested with 0.665μL of Shearase (Zymo) in a 10 μL volume at 42°C for 20min, inactivated at 65°C and purified with 0.8 volume of SPRI beads. End-repairing and A-tailing were performed by incubating in 25μL of repair mix (NEB End-repair buffer 2.5 μL, Klenow fragment with 3’-5’ exonuclease activity 0.5 μL, Taq polymerase 0.5 μL, T4 polynucleotide kinase 0.2 μL) at 25°C for 30min and 75°C for 30min, then purified with 1 volume of SPRI beads. Fragments were ligated to sequence adapters using quick ligase (NEB M2200S) in a total volume of 25 μL at 20°C for 20 min. Ligation mix was diluted to 50 μL with nuclease-free water, and purified by adding 29.15 μL SPRI beads and resuspended in 30 μL. PCR reaction was performed by adding 6μL of ligated fragments into 10 μL with Kapa Hifi HotStart ReadyMix and 2 μL of each barcoded primer (10mM), and cycling with the PCR program 98°C 40s, 9 cycles of 98°C 15s, 65°C 30s, 72°C 30s and finally 1 min at 72°C. After purifying with 0.8 volume of SPRI beads, samples were quantified with Qubit broad range DNA kit and pooled at equal ratios for sequencing on HiSeqX. SPRI beads were left in with the samples throughout the protocol, except before the quantification step during pooling.

For *Pachypanchax omalonotus* and *Nothobranchius virgatus*, a more simplified protocol was used. NEB Fragmentase (1 μL) was used to shear gDNA in 10 μL for 20min, stopped by 5μL of 0.5 M EDTA then purified with 1.2X SeraSure beads and eluted in 5 μL. End-repair and A-tailing were performed as before but in 10 μL, after which 15 μL of ligation mix was directly added to the uncleaned repair mix to a total volume of 25μL. Protocol then proceeded as before.

### Read processing and genotype calling

Paired-end reads were adapter- and quality-trimmed with Trimmomatic v 0.32 (Bolger, et al. 2014) using the following parameters: ILLUMINACLIP:adaptseqs.txt:3:7:7:1:true LEADING:10 TRAILING:10 SLIDINGWINDOW:4:10 MINLEN:20, and mapped using BWA-MEM (Li 2013) with default parameters to a version of the *P. playfairii* reference genome (La Digue) assigned by ALLMaps (Tang, et al. 2015) to pseudochromosomes by orthologous protein synteny information (Cui, et al. 2019) from *Xiphophorus maculatus* (weight = 100), *Nothobranchius furzeri* (weight = 80) and *Oryzias latipes* (weight=50).

PCR duplicates were marked with the MarkDuplicates tool from Picard tools v.1.119 (McKenna, et al. 2010) with default parameters. Cleaned reads were realigned around INDELs with the RealignerTargetCreator and IndelRealigner commands using default parameters in the GATK package (McKenna, et al. 2010). Variants were called by bcftools v1.6 with mpileup and call commands (Li, et al. 2009), with a minimal mapping quality of 30 for each individual. For McDonald Kreitman (MK) and Direction of Selection (DoS) calculations in the pool of *Pachypanchax* populations of Praslin, La Digue and Curieuse, variant calls were further converted to .pro format by Sam2Pro 0.6 with a minimal read coverage of 6. Allele frequencies were estimated by Package-GFE (Maruki and Lynch 2015) and polarized by using *Pachypanchax omalonotus*.

### Genome size estimation of *P. omalonotus*

As previously described, we mapped the trimmed reads of *P. omalonotus* to a draft genome of *P. playfairii* with BWA-MEM, counting read coverage in the single-copy BUSCOs. Genome size is calculated as total read bp divided by read coverage. This simple method was shown to agree well with flow-cytometry and not very sensitive to the choice of reference genomes or read coverage (Cui, et al. 2019).

### Phylogeny and molecular dating

To our previous dataset of orthologous protein-coding genes, we added newly available South American killifish genomes (Kelley, et al. 2016; Reid, et al. 2016; Wagner, et al. 2018) in addition to other representative teleost genomes from Ensembl and the newly obtained *Pachypanchx omalonotus*. Orthologs sequences were identified by the UPhO package (Ballesteros and Hormiga 2016), aligned, filtered and trimmed as previous described (Cui, et al. 2019). The resulting alignment contains 2,092,416 sites (697,472 codons) of 68 samples, including one individual from each *P. playfairii* population. The proportion of missing data ranges from 0-31.89% (mean 2.61%) per taxon. RAxML 8.2.9 (Stamatakis 2006) was run to estimate a phylogenetic tree using the GTR substitution model with gamma rate variation. Topological confidence was assessed by 100 rapid bootstraps. Before molecular dating, only a single sample was retained for each species in the dataset and the tree.

Divergence time was estimated by running MCMCTree (dos Reis and Yang 2011) using the GTR substitution model with a LogNormal distribution of clock rates, where clock rates were modeled as random variables for each branch independently. Four external calibration points were placed on the following nodes based on the posterior distributions in a previous study (Near, et al. 2012): MRCA(Spotted Gar, *Pachypanchax*) = 250 – 331.1 MYA, MRCA(Zebrafish, *Pachypanchax*) = 150.9–235MYA, MRCA(Takifugu, Tetraodon)=32–50MYA, MRCA(Medaka, *Pachypanchax*) = 49.1–130.8MYA. Two independent chains were run for 5 million generations with a sampling frequency of every 10 generations. The first 20,000 generations were discarded as burn-ins. Tracer was used to inspect the posterior traces of parameter estimates to ensure convergence between runs. The resulting tree was visualized with FigTree v1.4.4.

### Population genetic statistics

The gVCF files of individuals were merged with the merge command in bcftools v1.6, keeping missing genotypes as missing. The merged gVCF file was converted to .geno format with the parseVCF.py script from the genomic_general package, requiring a minimal variant quality of 60. We used the popgenWindows.py script (https://github.com/simonhmartin/genomics_general) to compute *F_ST_*, *d_xy_* and *θ_π_* for all *P. playfairii* populations in 20kb non-overlapping windows, with the requirement of at least 5000bp of non-missing data. The mean statistics across all windows are reported.

### TreeMix analysis

Distributions of read coverage at each population level were estimated from merged bcf files. BCF files were then converted into the TreeMix (Pickrell and Pritchard 2012) allele count format with a custom script, excluding sites with read coverage below the 0.025 or above the 0.975 coverage quantiles. The OptM R package (Fitak 2019) was used to determine the optimal number of migration edges. TreeMix was run with 0-6 migration edges in 20 iterations, starting from –k=200 to 4000, with an increment of 200 per iteration. The output files from TreeMix were used as input for OptM, where the Evano method was used to estimate the proportion of variance explained by different number of migration edges. The *ad hoc* statistics *Δm* was used to select for the optimal number of migration edges. We ran this analysis either with or leaving out the outgroup species *P. omalonotus*.

### PSMC analysis

We followed the MSMC2 (Schiffels and Durbin 2014) manual. Briefly, sequencing depth was estimated by averaging the per-base coverage computed by samtools depth command with no mapping quality cutoffs. The bamCaller.py tool from MSMC-Tool package was used to call variants and generate a mask of regions with abnormal coverage. The files were then converted to MSMC2 format by the generate_multihetsep.py tool. The PSMC’ method was run for each individual’s pseudo-chromosome separately using unphased genotypes in MSMC2, with an initial rho/mu set to 2.61976354 based on mu estimated from the dated phylogeny and the assumption of 2 recombinations/meiosis/chromosome. The mutation rate and recombination rate were then optimized by the program during the run.

### Asymptotic McDonald-Kreitman analysis and direction of selection

Calculations of the MK and DoS statistics are as previously described (Cui, et al. 2019).

### RELAX tests

Only single individuals were kept for RELAX (Wertheim, et al. 2014) and BUSTED-MH (Wisotsky, et al. 2020) tests. Pseudogenomes were made by aligning short reads to the closest related reference genomes as previously described. Protein-coding gene alignments were extracted, aligned and cleaned as described before. Due to the observation (simulation details at DOI: http://dx.doi.org/10.17632/f9phgbv4rs.1#file-0ec9fe0f-8085-44c5-9179-b9ee01c94360) that the original parameterization in the RELAX test would sometimes classify genes with both positive selection and relaxed selection as intensified (Cui, et al. 2019), resulting in missing some genes with a signature of relaxed selection, we modified the parameterization scheme to include two different **k** (intensification/relaxation) scaling parameters, where **k_1_** is used to transform site categories with ⍵ ≤ 1 (as ⍵^k^), and **k_2_** (also called q) applied to the site category with ⍵ > 1. Given two non-overlapping sets of tree branches (*reference* and *<u>test</u>*), the RELAX models describe selective regimes on the *reference* branches using a random effect with three ⍵ categories: ⍵_1_≤1 (weight p1), ⍵_2_≤1 (weight p2), and ⍵_3_≥1 (weight 1-p1-p2). Selective regimes on *test* branches follow the distribution ⍵_1_^k1^≤1 (weight p1), ⍵_2_^k1^≤1 (weight p2), and ⍵_3_ ^k2^≥1 (weight 1-p1-p2). Using this parametrization, we defined three models and fitted them to the data using maximum likelihood. Model 1 estimates **k_1_** and **k_2_** from the data, model 2 estimates **k_1_** but fixes **k_2_**=1 and model 3 fixes both parameters at 1. Relaxed purifying selection is tested by performing a likelihood ratio test between models 3 and 2 with 1 degree of freedom (**k_1_** > 1 implies intensification, and **k_1_** < 1 – relaxation). In theory, the comparison between Model 1 and Model 2 forms a test of positive selection, but simulations (**Table S2**) show that the power is very low and thus we do not rely on this test for positive selection (see below for BUSTED-MH instead). Compared to the original RELAX parameterization, the two-scaler parameterization has a reduced power to detect intensified purifying selection (17% of the original power, **Table S2**), thus we do not recommend it for that purpose. For our real dataset, we applied the two-scaler parameterization to detect relaxed genes, and the original one-scaler parameterization for detecting intensified genes. We provide the two-scaler parameterization analysis script (relax_q) for HYPHY 2.5.x (Pond, et al. 2019) in the supplementary information.

To compare the relative proportions of relaxed and intensified genes, we repeated these tests in a previously published dataset of continental African killifishes, with the addition the *Nothobranchius virgatus* genome.

### Positive selection test with BUSTED-MH

Detecting positive selection in protein-coding sequences has recently been shown to be vulnerable to two violations in model assumptions, namely instantaneous multinucleotide substitutions (Venkat, et al. 2018) and site-to-site variation in synonymous substitution rates (Wisotsky, et al. 2020). We thus use a developmental version of the BUSTED test that accounts for both processes, named BUSTED-MH, in order to test for positive selection in the genus *Pachypanchax*. This script implementing the analysis can be run with HYPHY v2.5.10 or later and is available from https://github.com/veg/hyphy-analyses/.

### Conserved element detection and PhastCon analysis

We performed a whole-genome alignment using Progressive Cactus (Paten, et al. 2011) of 4 species in three annual genera (*Austrofundulus*, *Callopanchax* and *Nothobranchius*), and 1 non annual species (*P. playfairii*). PhastCons in the R rphast package (Hubisz, et al. 2011) was used to identify conserved genomic regions in the annual species, leaving out the non-annual. Due to the long divergence of these annual genera, the conserved genomic regions likely contain functionally important regulatory or protein-coding elements. After excluding protein-coding elements, we then identified accelerated non-coding regions in *P. playfairii* that are otherwise conserved in annuals by comparing the rate with 4-fold degenerate sites with PhyloFit in the rphast package. Because 4-fold degenerate sites evolve slower than neutrality in many other organisms (Künstner, et al. 2011; Lawrie, et al. 2013), accelerated conserved elements (ACEs) detected in this manner likely reflect relaxed selection.

### DREME analysis

We extracted the 50bp *Pachypanchax*-ACE sequences from the homologous locations in *N. furzeri*, and used DREME (Bailey 2011) to analyze the enrichment of hexamers. A literature search was performed to check whether any of the enriched hexamers corresponds to known functional regulatory motifs.

### Gene ontology and pathway analysis

Gene ontology and pathway enrichment analyses were performed on the ConsensusPathDB website (Herwig, et al. 2016). Genes called relaxed (p < 0.05) were mapped to Ensembl IDs of the human ortholog, and compared to a background list of all genes that were entered into the RELAX analysis. For the ACEs, the immediately down-stream protein-coding gene was used as the foreground, and a random set of genes sampled from the genome which are more than 170kb away from any ACEs was used as the background.

### Incubation of killifish embryos in sea water

We directly compared viability of killifish embryos incubated in autoclaved fish room water and artificial sea water (33 g RedSea Marine Salt in 1L RO water). Four species/strains were examined: *P. playfairii*, *N. furzeri* GRZ-AD, GRZ-Bell and MZCS-002. For each species or strain, fertilized eggs produced within 1 week were collected from 3-4 breeding tanks each consisted of 1 male and 3-4 females. Eggs in batch 1 was bleached with 0.5% hydrogen peroxide to prevent fungal infections. We used n=100 eggs per *N. furzeri* strain and n=117 eggs for *P. plaifairii*, with approximately 25-30 eggs per petri dish. Batch 2 was washed in autoclaved fresh water but not bleached. Batch 2 contains 320 GRZ-Bell eggs, 140 MZCS-002 eggs and 94 *P. playfairii* eggs, incubated at a density of 23-40 eggs/petri dish. In both batches, embryos were checked daily. Dead embryos were recorded, removed and medium was exchanged. Hatched *P. playfairii* were transferred to the hatching incubator and slowly acclimatized to freshwater conditions if they were hatched in sea water (by replacing 50% of the artificial sea water with freshwater daily). *N. furzeri* embryos were transferred to damp Whatman paper after reaching black eye stage for incubation. All embryos were incubated at 28°C.

## Acknowledgments

We thank Francesca and Jacopo Valdesalici for their help on field sampling, the residents of Seychelles for their continuous support and kindness. Hilton Seychelles Labriz Resort & Spa provided logistics and transport to Silhouette island. We would like to thank Francois Baguette and Ella (Silhouette Island Conservation Officer) for help in sampling fish on Silhouette island. We thank the members of the Valenzano lab at the Max Planck Institute for Biology of Ageing for continuous discussions and feedback. All computations were performed on the Amalia HPC cluster at MPI-AGE. This work was funded by the Max Planck Institute for Biology of Ageing and by the Max Planck Society.

## Author contributions

D.R.V., S.V., R.C. conceived and designed the study. S.V. and E.H. collected fish samples. Z.M and R.C. made the sequencing libraries. A.T. performed sea-water embryo incubation. S.P. and R.C. implemented the 2-scaler RELAX test and BUSTED-MH. R.C. analyzed the data. R.C. and D.R.V. wrote the manuscript. All authors read and commented the manuscript.

---

**Figure S1.**
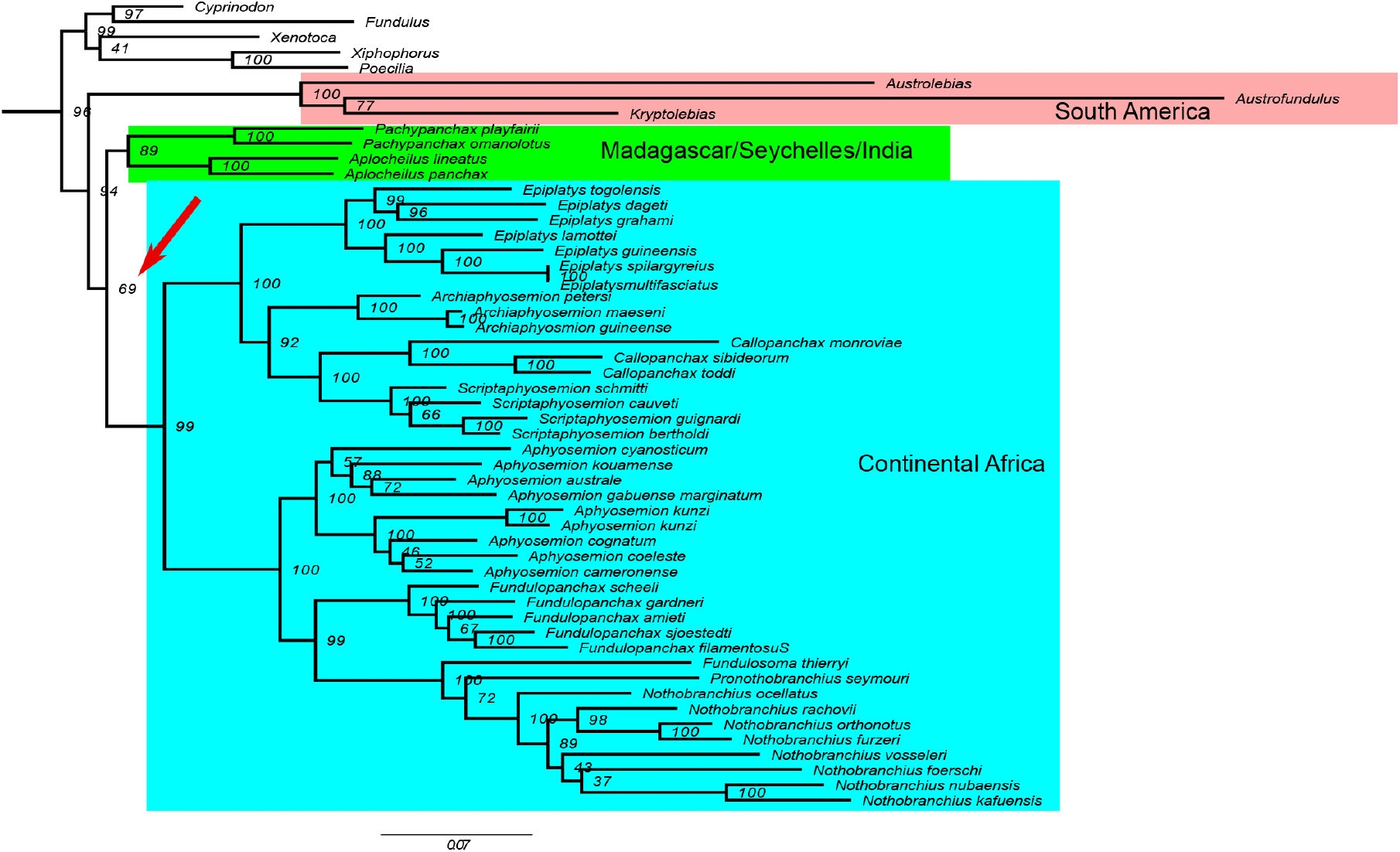
Maximum likelihood tree inferred by RAxML on codon positions 1 and 2 of all mitochondrial protein-coding genes. AU test does not reject the alternative topology of monophyly between South American and African taxa, but the ML tree agrees with the nuclear genome.

**Figure S2.**
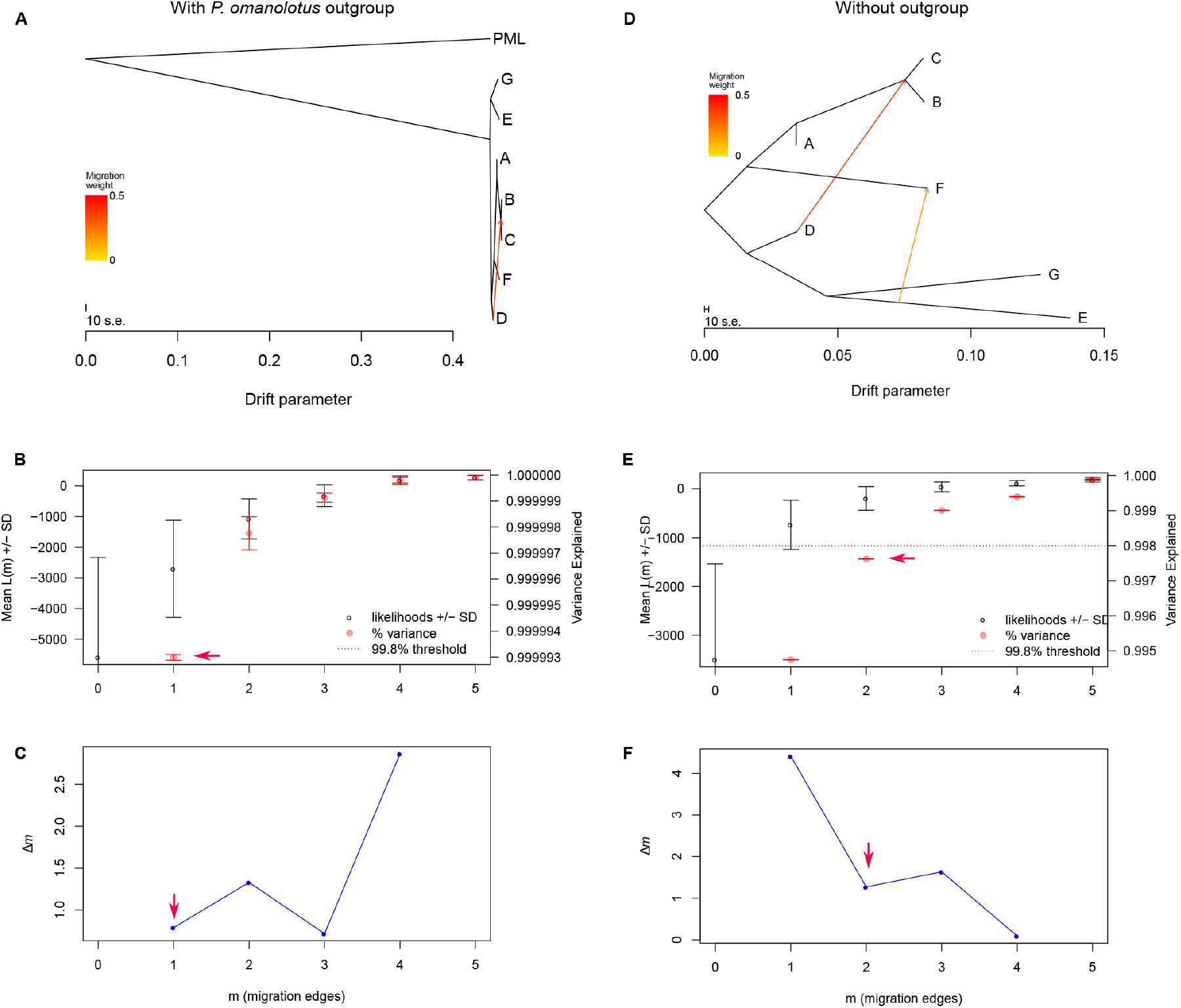
TreeMix analyses with (**A-C**) or without (**D-F**) *P. omalonotus* as an outgroup. **A, B** The TreeMix graph with the optimal number of migration edges identified by OptM. k=400 was used for plotting. **B, E.** Selection of the most likely number of migration edges by the OptM R package. Distribution of log likelihood and variance explained of Treemix models with 0-5 migration edges. Standard deviation generated by repeating analyses by varying k (snps per window) from 200 to 4000, with a 200 increment. The number of migration edges that explains ~99.8% of the variance is chosen as the optimal number of edges. **C. F.** Selection of the most likely number of migration edges by plotting deltaM (OptM package).

**Figure S3.**
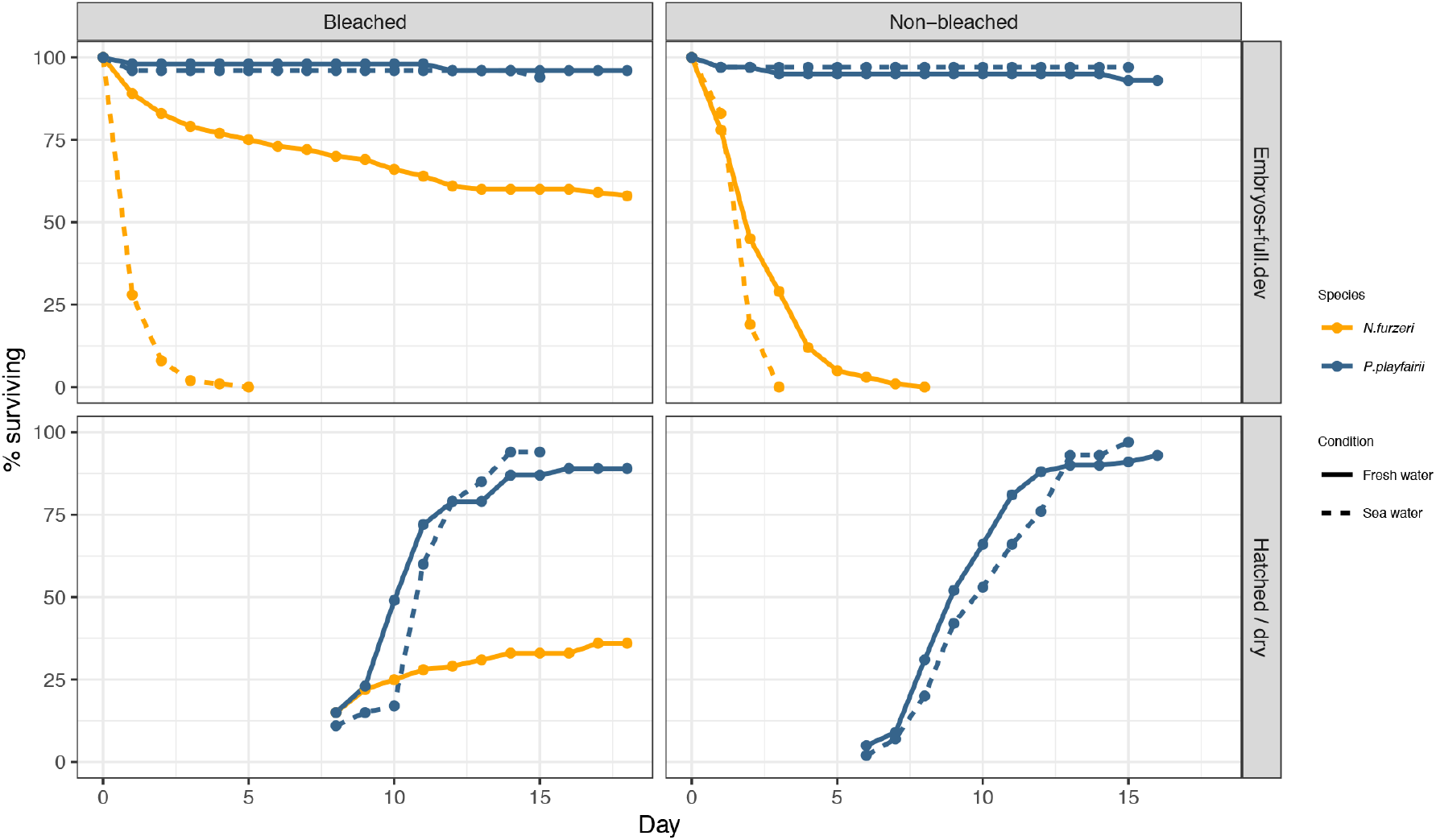
Survival rate of *P. playfairii* (blue) and *N. furzeri* (orange) embryos in freshwater and sea water. Left column shows incubation of eggs bleached with 0.5% hydrogen peroxide prior to experiment for fungal removal. Right column shows incubation of freshly collected eggs without fungal removal. Top row shows total embryo survival (including successfully hatched) rates of the two killifish species in freshwater (solid line) and sea-water (dotted line). Bottom row shows hatched and surviving fry.

**Figure S4.**
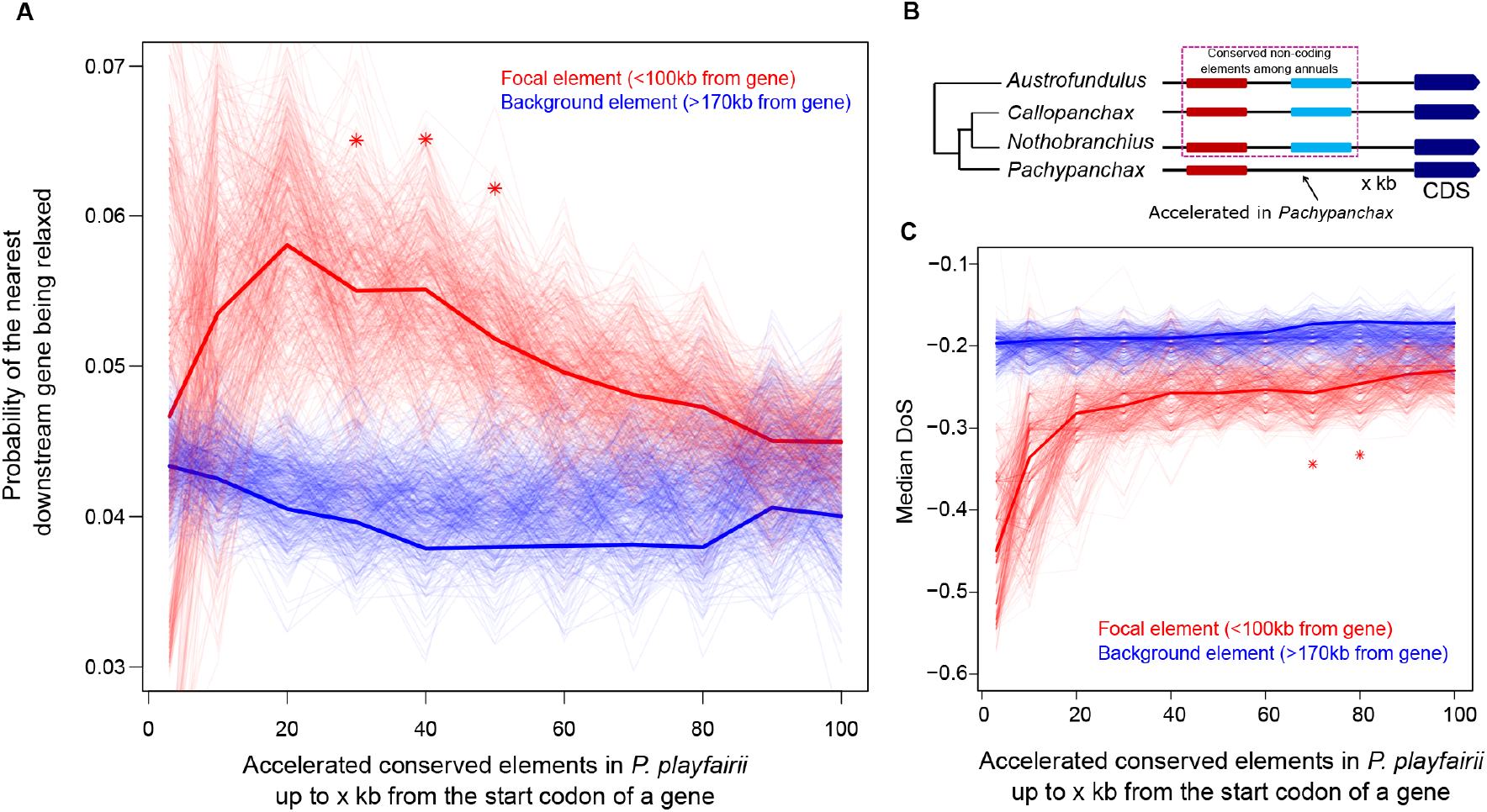
**A.** The probability of a gene immediately downstream of an accelerated conserved element in *P. playfairii* to be relaxed in the lineage *Pachypanchax* is higher. Asterisks mark Fisher's exact tests with p < 0.05. Faint lines generated by 500 bootstraps. **B.** An ideogram showing the method for detecting accelerated conserved elements in the *Pachypanchax* genome. The protein coding gene immediately downstream of the focal element was used for correlation. **C.** The median direction of selection (DoS) of the protein coding genes immediately downstream of accelerated conserved elements in *P. playfairii* is more negative. Asterisks mark Wilcoxon rank sum tests with p < 0.05. Faint lines generated by 500 bootstraps.

**Figure S5.**
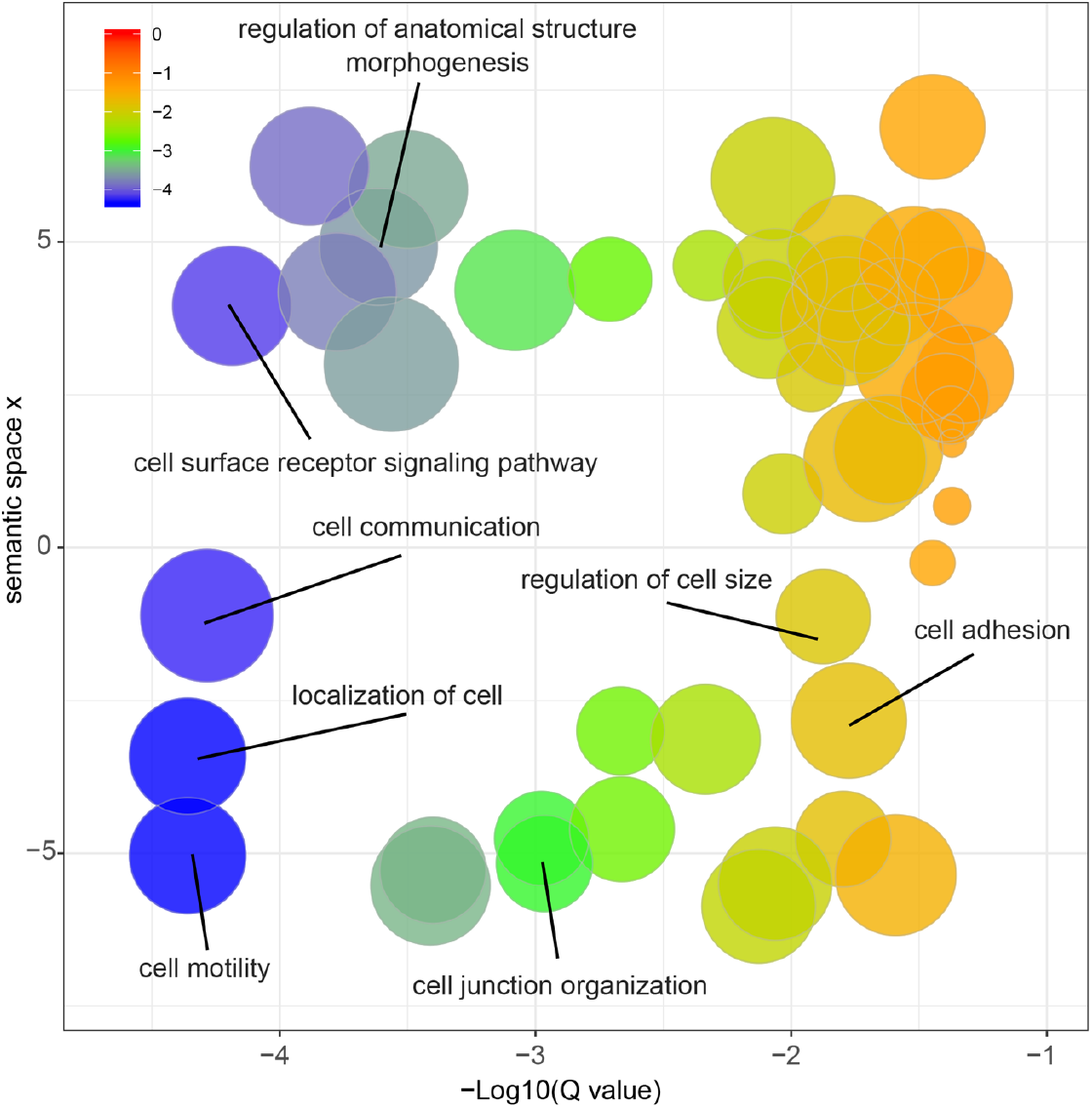
Enriched gene ontology terms (q < 0.05) of the relaxed genes (p<0.05, k<1) detected in the *Pachypanchax* lineage plotted on a semantic space using Evigo.

**Figure S6.**
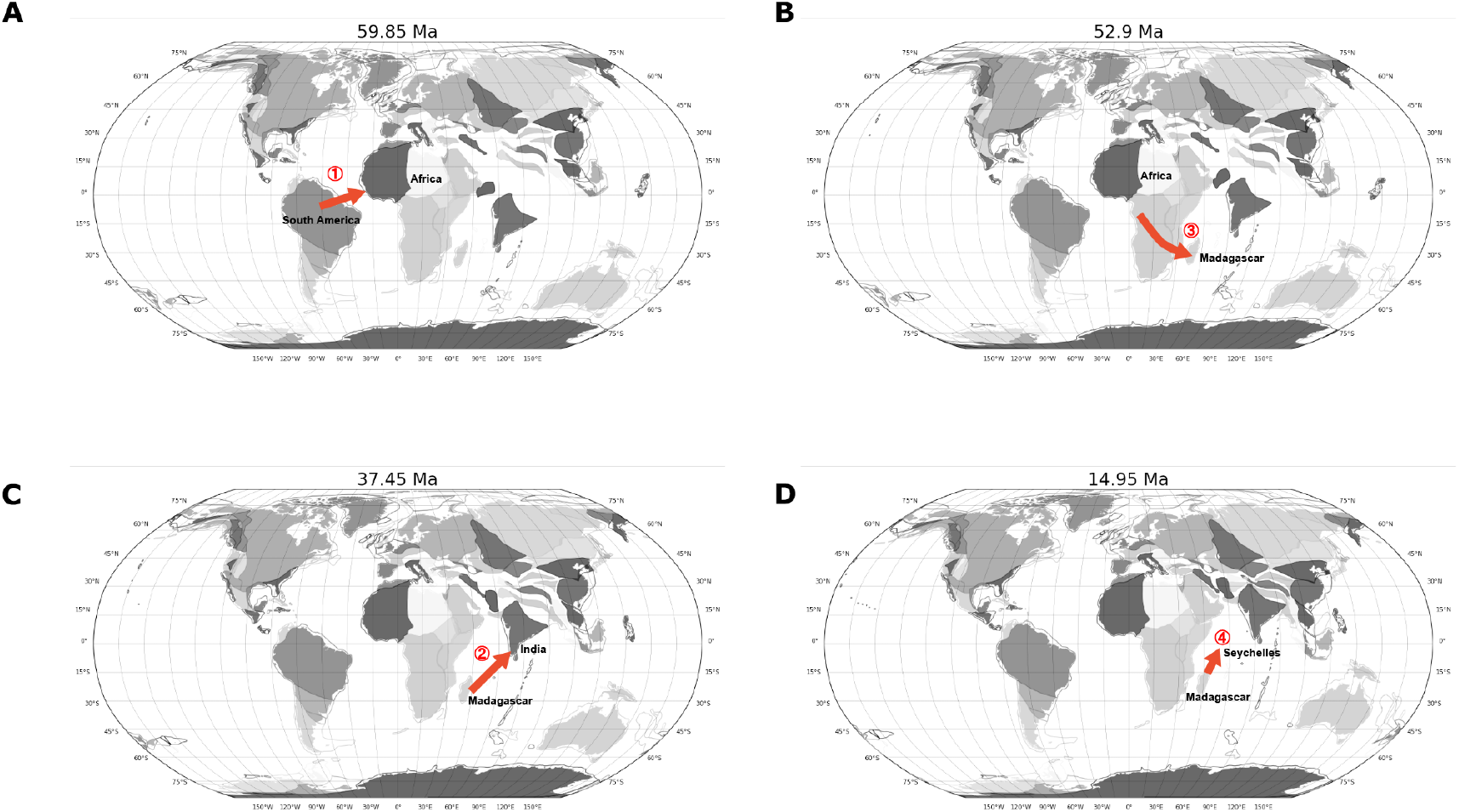
Putative transoceanic dispersal events plotted on reconstructed PALAEOMAP (https://www.earthbyte.org/paleomap-paleoatlas-for-gplates/) at the inferred divergence times of killifish. The speciation event numbering follows **Figure 1**.

